# Meningeal lymphatic dysfunction exacerbates traumatic brain injury pathogenesis

**DOI:** 10.1101/817023

**Authors:** Ashley C. Bolte, Mariah E. Hurt, Igor Smirnov, Michael A. Kovacs, Celia A. McKee, Nick Natale, Hannah E. Ennerfelt, Elizabeth L. Frost, Catherine E. Lammert, Jonathan Kipnis, John R. Lukens

**Author notes:** **Correspondence should be addressed to:** John R. Lukens, Department of Neuroscience, Center for Brain Immunology and Glia, University of Virginia, 409 Lane Road, MR4- 6154, Charlottesville VA 22908.

## Abstract

Traumatic brain injury (TBI) has emerged as a leading cause of death and disability. Despite being a growing medical issue, the biological factors that promote central nervous system (CNS) pathology and neurological dysfunction following TBI remain poorly characterized. Recently, the meningeal lymphatic system was identified as a critical mediator of drainage from the CNS. In comparison to other peripheral organs, our understanding of how defects in lymphatic drainage from the CNS contribute to disease is limited. It is still unknown how TBI impacts meningeal lymphatic function and whether disruptions in this drainage pathway are involved in driving TBI pathogenesis. Here we demonstrate that even mild forms of brain trauma cause severe deficits in meningeal lymphatic drainage that can last out to at least two weeks post-injury. To investigate a mechanism behind impaired lymphatic function in TBI, we examined how increased intracranial pressure (ICP) influences the meningeal lymphatics, as increased ICP commonly occurs in TBI. We demonstrate that increased ICP is capable of provoking meningeal lymphatic dysfunction. Moreover, we show that pre-existing lymphatic dysfunction mediated by targeted photoablation before TBI leads to increased neuroinflammation and cognitive deficits. These findings provide new insights into both the causes and consequences of meningeal lymphatic dysfunction in TBI and suggest that therapeutics targeting the meningeal lymphatic system may offer strategies to treat TBI.

## INTRODUCTION

Traumatic brain injury (TBI) affects millions of people worldwide each year, and current estimates from the World Health Organization suggest that TBI will be the third leading cause of death and disability by the year 2020 [1]. TBI can cause debilitating impairments in motor function, cognition, sensory function, and mental health. In addition, mounting evidence indicates that having a history of TBI markedly increases the risk of developing numerous other neurological disorders later in life including chronic traumatic encephalopathy (CTE), Alzheimer’s disease, anxiety, depression, and amyotrophic lateral sclerosis [2-7]. Despite being a prevalent and pressing global medical issue, the pathoetiology of TBI remains incompletely understood and improved treatment options are desperately needed.

TBI results in damage to the brain through death of CNS resident cells including neurons, glia and meningeal cells. This tissue damage and cellular stress promote immune responses that are intended to aid in the disposal of neurotoxic material and coordinate tissue repair [8, 9]. While immune responses initially play beneficial roles in TBI, unchecked and/or chronic immune activation following brain trauma can lead to secondary tissue damage, brain atrophy, and eventual neurological dysfunction [8-13]. Notably, the inability to properly dispose of danger/damage-associated molecular patterns (DAMPs) such as protein aggregates, necrotic cells, and cellular debris has been shown to be a pivotal driver of both persistent and maladaptive immune activation in numerous neurological disorders [14, 15]. In the case of CNS injury, inefficient removal of DAMPs has been proposed to perpetuate neuroinflammation and incite secondary CNS pathology and neurological complications [9, 14, 16, 17]. However, we currently lack complete knowledge of the drainage pathways that the brain relies on to dispose of DAMPs and resolve tissue damage following TBI.

Emerging studies over the last few years have shown that the meningeal lymphatics are centrally involved in the drainage of macromolecules, cellular debris, and immune cells from the brain to the periphery during homeostasis [18, 19]. The anatomy and function of this CNS drainage pathway are just now being defined, and its role in many neurological diseases, including TBI, has not been elucidated [18-23]. In recently published work, it was shown that these lymphatic vessels drain cerebrospinal fluid (CSF), interstitial fluid (ISF), CNS-derived molecules, and immune cells from the brain and meninges to the deep cervical lymph nodes (dCLN) [18, 19, 22]. Importantly, studies using *in vivo* magnetic resonance imaging (MRI) techniques have also identified the existence of meningeal lymphatic vessels in both humans and nonhuman primates [24, 25]. More recent studies have also shown that the meningeal lymphatic system is critical for clearing amyloid beta, extracellular tau, and alpha synuclein from the brain, and that disruption of this drainage system can promote the accumulation of these neurotoxic DAMPs in the brain [21, 26, 27]. Whether meningeal lymphatic dysfunction plays a role in TBI currently remains poorly understood.

Here, we explored how the meningeal lymphatics are impacted following TBI, and how possessing defects in this drainage system before brain trauma influences TBI pathogenesis. We find that TBI results in compromised meningeal lymphatic drainage that can last out to 2 weeks post-injury. We also report that pre-existing deficits in meningeal lymphatic function predispose the brain to exacerbated clinical disease following brain trauma. Moreover, we shed light on a mechanism to explain why TBI results in CNS lymphatic dysfunction by highlighting that increased intracranial pressure (ICP) is capable of promoting impaired meningeal lymphatic drainage.

## METHODS

### Mice

All mouse experiments were performed in accordance with the relevant guidelines and regulations of the University of Virginia and approved by the University of Virginia Animal Care and Use Committee. C57BL/6J mice were obtained from The Jackson Laboratories. Mice were housed and behavior was conducted in specific pathogen-free conditions under standard 12-h light/dark cycle conditions in rooms equipped with control for temperature (21 ± 1.5°C) and humidity (50 ± 10%). Mice matched for sex and age were randomly assigned into experimental groups.

### Traumatic Brain Injury

Mice were anesthetized by 4% isoflurane with 0.3kPa O_2_ for 2 mins and then the right preauricular area was shaved. The mouse was placed prone on a foam bed with its nose secured in a nosecone delivering 1.5% isoflurane. The device used to deliver TBI was a Controlled Cortical Impact Device (Leica Biosystems, 39463920). A 3mm impact probe was attached to the impactor device which was secured to a stereotaxic frame and positioned at 45 degrees from vertical. In this study, we used a strike depth of 2 mm, 0.1 secs of contact time and an impact velocity of 5.2m/s. The impactor was positioned at the posterior corner of the eye, moved 3 mm towards the ear and adjusted to the specified depth using the stereotaxic frame. A cotton swab was used to apply water to the injury site and the tail in order to establish contact sensing. To induce TBI, the impactor was retracted and dispensed once correctly positioned. Following impact, the mouse was placed supine on a heating pad and allowed to regain consciousness. After anesthesia induction, the delivery of the injuries took less than 1 minute. The time until the mouse returned to the prone position was recorded as the righting time. Upon resuming the prone position, mice were returned to their home cages to recover on a heating pad.

### Intra-cisterna magna injections

Mice were anaesthetized by intraperitoneal (i.p.) injection of a mixed solution of ketamine (100 mg/kg) and xylazine (10 mg/kg) in sterile saline. The skin of the neck was shaved and cleaned with iodine and 70% ethanol, and ophthalmic solution (Puralube Vet Ointment, Dechra) was placed on the eyes to prevent drying. The head of the mouse was secured in a stereotaxic frame and an incision in the skin was made at midline. The muscle layers were retracted and the cisterna magna exposed. Using a Hamilton syringe (coupled to a 33-gauge needle), the volume of the desired solution was injected into the cerebrospinal fluid (CSF)-filled cisterna magna compartment. For the bead experiments, 2 μl of FluoSpheres carboxylate 0.5μm-505/515 (Invitrogen) in artificial CSF (597316, Harvard Apparatus UK) were injected at a rate of 2μl/min. For Visudyne experiments, 5 μl of Visudyne (verteporforin for injection, Valeant Opthalmics) was injected at a rate of 2.5 μl/min. For Lyve-1 labeling experiments, 2 μl of anti-mouse Lyve1-488 (Invitrogen, 53044382, undiluted) was injected at a rate of 2 μl/min. The needle was inserted into the cisterna magna through retracted muscle in order to prevent backflow upon needle removal. The neck skin was then sutured, after which the mice were subcutaneously injected with ketoprofen (1 mg/kg) and allowed to recover on a heating pad until fully awake.

### Pharmacologic meningeal lymphatic vessel ablation

Visudyne treatment was adapted from previously published protocols [20, 21, 28]. Selective ablation of the meningeal lymphatic vessels was achieved by i.c.m. injection and transcranial photoconversion of visudyne (verteporfin for injection, Valeant Ophthalmics). Visudyne was reconstituted following the manufacturer’s instructions and 5 μl was injected i.c.m. following the procedure described above in ‘Intra-cisterna magna injections’. After 15 min, a midline incision was created in the skin to expose the skull bone and visudyne was photoconverted by pointing a 689-nm-wavelength non-thermal red light (Coherent Opal Photoactivator, Lumenis) to five different locations above the intact skull (1 at the injection site, 1 at the superior sagittal sinus, 1 at the confluence of the sinuses and 2 at the transverse sinuses). This experimental group is labeled as ‘Visudyne + laser’ or ‘Ablated’. Each location was irradiated with a light dose of 50 J/cm^2^ at an intensity of 600 mW/cm^2^ for a total of 83 sec. Controls were injected with the same volume of visudyne (without the photoconversion step; labeled as ‘Visudyne’) or sterile saline plus laser treatment (labeled as ‘Vehicle + laser’). The scalp skin was then sutured, after which the mice were subcutaneously injected with ketoprofen (1 mg/kg) and allowed to recover on a heating pad until fully awake.

### Intracranial pressure measurements

Intracranial pressure (ICP) was measured as previously described [20, 21]. Mice were anaesthetized by i.p. injection with ketamine (100 mg/kg) and xylazine (10 mg/kg) in saline and the skin was incised to expose the skull. A 0.5-mm diameter hole was drilled in the skull above the left parietal lobe. Using a stereotaxic frame, a pressure sensor catheter (model SPR100, Millar) was inserted perpendicularly into the cortex at a depth of 1 mm. To record changes in ICP, the pressure sensor was connected to the PCU-2000 pressure control unit (Millar). For measurements in mice after TBI (30min, 2hr, 6hr, 24hr, 3day, 4day and 1wk post injury) or after jugular venous ligation (2hr and 24hr) ICP was recorded for 6 min after stabilization of the signal and the average pressure was calculated over the last 3 min of recording. Mice were euthanized following the procedure.

### Jugular venous ligation

Mice were anaesthetized by i.p. injection with ketamine (100 mg/kg) and xylazine (10 mg/kg) in saline. The left and right preauricular area and the skin between the ears was shaved and prepped with iodine and 70% ethanol. Ophthalmic ointment (Puralube Vet Ointment, Dechra) was applied to the eyes to prevent drying. The mouse was secured onto a surgical plane in the lateral position and a lateral incision was created between the two mouse ears. The incision site was retracted to reveal the left temporalis muscle. The left temporalis muscle was retracted to reveal the infratemporal fossa, where the left internal jugular vein can be identified. The left internal jugular vein was ligated using 8-0 Nylon Suture (AD surgical, XXS-N808T6), and then the same procedure was performed on the opposite side to ligate the right jugular vein. The incision was then sutured and the mice were subcutaneously injected with ketoprofen (1 mg/kg) and were allowed to recover on the heating pad until awake. Sham mice received the incision and the jugular veins were exposed bilaterally, but they did not undergo ligation. The intracranial pressure on these mice was recorded as described in ‘intracranial pressure measurements’ 3 and 24 hrs after ligation.

### Behavioral Testing

All behavioral experiments were carried out during daylight hrs (except the Novel Object Recognition Test, which was carried out starting at 7:00PM) in a blinded fashion.

### Gross Neuroscore

The gross neuroscore was performed following a published protocol with modifications [29]. Briefly, mice performed 10 individual tasks to assess behaviors including seeking/exploring tendencies, the startle reflex, and balance/motor coordination. The ability to cross different width beams, to react to a loud noise, to balance on a beam and to explore the surroundings were assessed and scored by a blinded experimenter 1 hr after TBI. If the mouse was able to adequately perform the task, a score of 0 was given. If the mouse failed to adequately perform the test, a score of 1 was given. Scores for the 10 tasks were summed for a total minimum score of 0 and a total maximum score of 10.

### Rotarod test

The mice were transported to the behavior room and allowed to habituate for 1 hour before each day of testing. The experimental apparatus used in this test contained 5 separate compartments on a rotating rod to accommodate 5 mice per trial (MED Associates Inc, ENV-575M). The rod was programmed to turn starting at 4 rotations per minute (rpm) and to accelerate to 40 rpm through a span of 5 mins. Each mouse was placed on the rod and allowed to ambulate until it either fell off, hung without effort on the rod for a total of 5 rotations, or reached the trial endpoint (6 min). When a mouse falls from the rotarod, it disrupts a laser sensor to stop recording. The time spent on the rod and the speed at which the mouse fell or the trial ended was recorded (RotaRod Version 1.4.1, MED associates inc). Three trials were performed each day for 3 days. The three trials per day were averaged, and latency to fall and percent performance increase were calculated based off of the average time of trial per mouse per day.

### Novel object recognition test (NORT)

The novel object recognition test was performed following a published protocol with modifications [30]. The mice were transported to the behavior room and allowed to habituate for 1 hour before each trial of the test. The experimental apparatus used in this study was a square box made of opaque white plastic (35 cm × 35 cm). The mice were first habituated to the square apparatus for 10 min by allowing for free exploration within the open field. 10 hrs later, after 7:00PM, two identical objects (both red with identical shapes and textures) were then positioned in the two far corners of the arena at distances of 5 cm away from the adjacent arena wall (familiar objects). Mice were then placed in the arena facing the wall furthest away from the objects and allowed to explore the arena and objects for 10 min. Time spent investigating the objects was measured and was considered the “training phase” of the test. After 24 h, the mice were placed in the same box with two objects in the same locations, but one of the familiar objects was exchanged with a novel object that had a different shape, texture and color but similar dimensions as the original object (novel object). The time spent exploring the familiar and novel objects was measured for 10 min and was considered the “test phase”. Exploration of an object was recorded when the mouse approached an object and touched it with its vibrissae, snout or forepaws and was measured using a video tracking software (TopScan, CleverSys, Inc.). The preference for either the novel or familiar object was calculated based on the time each mouse spent investigating the two objects.

### Morris water maze (MWM) test

The MWM test was performed as previously described [21, 31], but with minor modifications. Mice were transported to the behavior room to habituate at least 1 hour before starting the test each day. The MWM test consisted of four days of training, one day of probe trial and two days of reversal. In the training, mice performed 4 trials per day, for 4 consecutive days, to find a hidden 10-cm diameter platform located 1 cm below the water surface in a pool that was 1 m in diameter. Tap water was made opaque with nontoxic tempera white paint (PRANG, 10607) and the water temperature was kept at 23 ± 1 °C. A dim light source was placed within the testing room and distal visual cues were available above each quadrant of the water maze to aid in spatial navigation and location determination for the submerged platform. The latency to platform, the time required by the mouse to find and climb onto the platform, was recorded for up to 60 s. After the first trial, each mouse was allowed to remain on the platform for 2 min to allow the mouse to observe its surroundings and then was moved from the maze to its home cage. If the mouse did not find the platform within 60 secs, it was manually placed on the platform and returned to its home cage after 2 min. After the first trial, the mouse was left on the platform for 10 secs. The inter-trial interval for each mouse was at least 30 min. On day 5, the platform was removed from the pool, and each mouse was tested in a probe trial for 60 secs. On days 1 and 2 of the reversal trial phase, without changing the position of the visual cues, the platform was placed in the quadrant opposite to the original acquisition quadrant and the mouse was retrained for four trials per day. All MWM testing was performed during the lights-on phase, by a blinded experimenter. During the training, probe and reversal tests, data was recorded using an automated tracking system (Noldus Information Technology, Version 14.0). The mean latency (in secs) of the four trials was calculated for each day of test trials. The percentage of time in the platform quadrant was calculated for the probe trial.

### Tissue collection

Mice were euthanized with CO_2_ and then transcardially perfused with 20mL PBS. Deep cervical lymph nodes were dissected and drop-fixed in 4% paraformaldehyde (PFA) for 2 hr at 4 °C and then the CUBIC clearance protocol was performed as previously described [32]. For meningeal whole mount collection, skin and muscle were stripped from the outer skull and the skullcap was removed with surgical scissors and fixed in 2% PFA for 12 hrs at 4 °C. Then the meninges (dura mater and arachnoid mater) were carefully dissected from the skullcaps with Dumont #5 forceps (Fine Science Tools). Meningeal whole-mounts were then moved to PBS and 0.05% azide at 4 °C until further use. Brains were removed and kept in 4% PFA for 24 hr, cryoprotected with 30% sucrose for 3 days, and frozen in Tissue-Plus OCT compound (Thermo Fisher Scientific). Fixed and frozen brains were sliced (50-μm thick sections) with a cryostat (Leica) and kept in PBS and 0.05% azide at 4 °C until further use.

### Immunohistochemistry, imaging and quantification

For immunofluorescence staining, floating brain sections and meningeal whole-mounts in PBS and 0.05% azide were blocked with either 2% donkey serum or 2% Goat serum, 1% bovine serum albumin, 0.1% triton, 0.05% tween, 0.05% sodium azide in PBS for 1.5 hr at room temperature. This blocking step was followed by incubation with appropriate dilutions of primary antibodies: anti-LYVE-1–eFluor 660 & eFluor 488 (eBioscience, clone ALY7, 1:200), anti-CD31 (Millipore Sigma, MAB1398Z, clone 2H8, 1:200), anti-Iba1 (Abcam, ab5076, 1:300) and anti-GFAP (Thermo Fisher Scientific, 2.2B10, 1:1000) in the same solution used for blocking overnight at 4°C or for 3 hrs at RT. Meningeal whole-mounts or brain tissue sections were then washed three times for 10 min at room temperature in PBS and 0.05% tween-20, followed by incubation with the appropriate goat or donkey Alexa Fluor 488, 546, 594 or 647 anti-rat, -goat, -rabbit, -mouse (Thermo Fisher Scientific, 1:1000) or -Armenian hamster (Jackson ImmunoResearch, 1:1000) IgG antibodies for 2 hrs at RT in the same solution used for blocking. The sections or whole-mounts were then washed 3 times for 10 mins at RT before incubation for 10 min with 1:1000 DAPI in PBS. The tissue was then transferred to PBS and mounted with ProLong Gold antifade reagent (Invitrogen, P36930) on glass slides with coverslips. Slide preparations were stored at 4°C and imaged using a Lecia TCS SP8 confocal microscope and LAS AF software (Leica Microsystems) within one week of staining. Quantitative analysis of the acquired images was performed using Fiji software. For the assessment of gliosis in the hemisphere affected by the injury, two representative slides of the brain sections were imaged and the mean area fraction was calculated using Microsoft Excel. For lymph nodes, the % volume of microbead coverage in cleared dCLN was assessed by creating a 3D reconstruction of the node and calculating the volume covered by beads divided by the total volume of the node using Fiji. The right and the left dCLN % volume were averaged together for each mouse. For assessment of meningeal lymphatic vessel coverage and complexity, images of meningeal whole-mounts were acquired using a confocal microscope and Fiji was used for quantifications. When applicable, the same images were used to assess the percentage of field coverage by LYVE-1^-^ CD31^+^ vessels.

### Statistical analysis and reproducibility

Sample sizes were chosen on the basis of standard power calculations (with *α* = 0.05 and power of 0.8). Experimenters were blinded to the identity of experimental groups from the time of euthanasia until the end of data collection and analysis. One-way ANOVA, with Bonferroni’s post hoc test or Holm-Sidak’s post hoc test, was used to compare three independent groups. Two-group comparisons were made using unpaired Students t-test. For comparisons of multiple factors (for example, age versus treatment), two-way ANOVA with Bonferroni’s post hoc test was used. Repeated-measures two-way ANOVA with Bonferroni’s post hoc test was used for day versus treatment comparisons with repeated observations. Statistical analysis (data are always presented as mean ± s.e.m.) was performed using Prism 8.0 (GraphPad Software, Inc.).

## RESULTS

### TBI causes meningeal lymphatic dysfunction

To investigate whether meningeal lymphatic drainage function is impacted by TBI, we employed a mild closed-skull model of TBI that uses a stereotaxic electromagnetic impactor to deliver a single blow to the right inferior temporal lobe. This model is ideal for studying CNS lymphatic function because it does not rely on a craniotomy to perform the TBI and also does not result in a direct impact to the lymphatic vessels (Figure 1a). Following TBI induction, injured mice had an average righting time of 300 seconds post-injury (Figure 1b) and TBI mice performed as well as sham mice in a series of behavioral tasks assessing injury-associated deficits in balance, motor coordination, reflex, and alertness (Figure 1c). In addition, brain injury in this model did not affect performance on an accelerating rotarod (Figure 1d) and brains from TBI mice only showed modest increases in Iba-1 and GFAP staining (Supplementary Figure 1).

**Figure 1.**
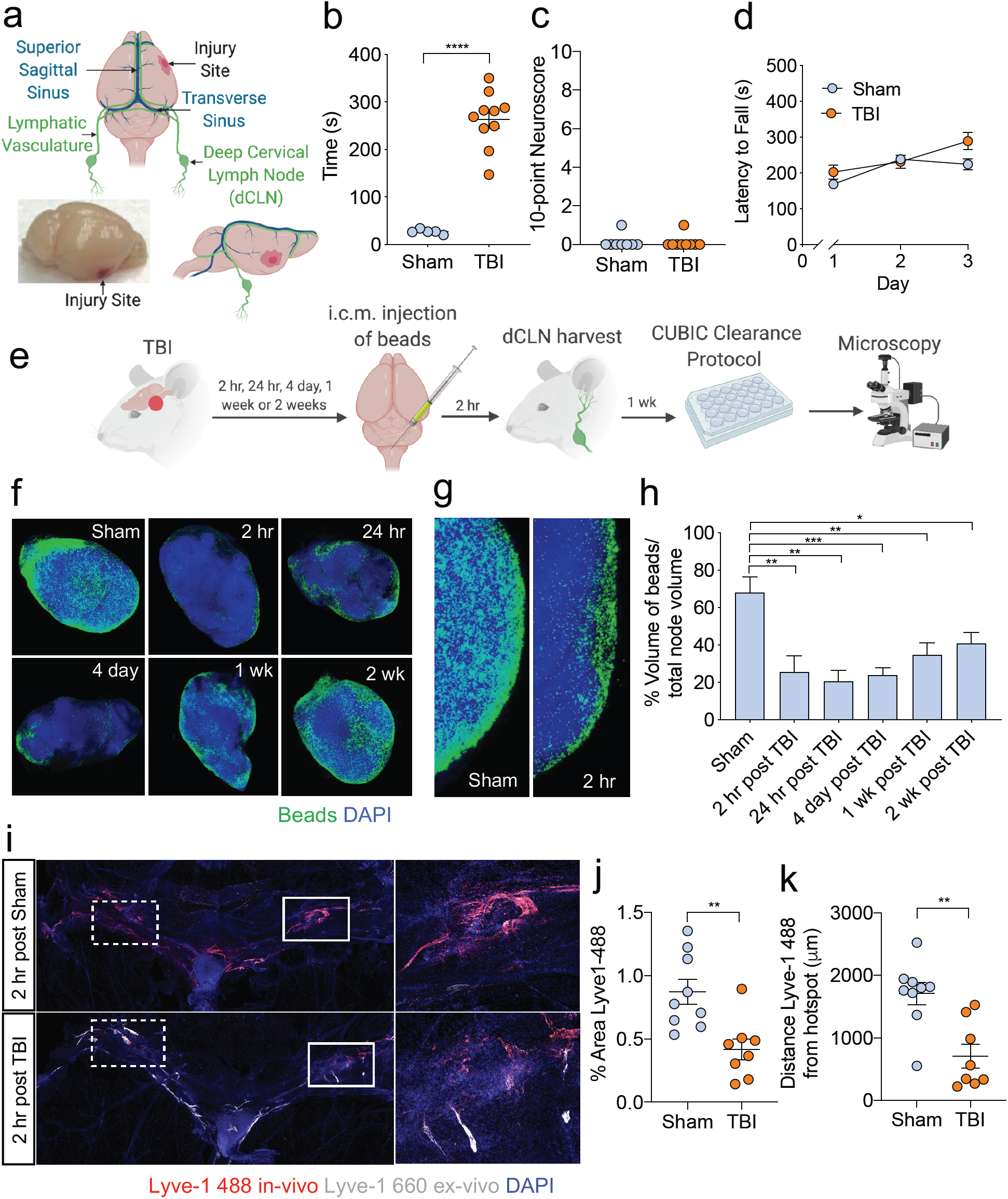
TBI leads to impairments in meningeal lymphatic drainage. a) Location of injury site in relation to the CNS lymphatic vasculature. b) Righting times of TBI and sham mice, representative data from 10 independent experiments. c) The 10-point gross neuroscore test was used to assess neurological function 1 hr after TBI, representative data from two independent experiments. d) The accelerating rotarod behavioral test was used to assess motor function the first three days after TBI, representative data from two independent experiments, n=10 mice/group. e) Schematic of the experimental layout where mice received TBI and then were injected i.c.m. with 0.5 μm fluorescent beads. dCLN were harvested and cleared according to the CUBIC protocol. (f,g) Representative images and (h) quantification of bead accumulation in the cleared dCLN at 2 hr, 24 hr, 4 day, 1 wk and 2 wk after TBI. Each data point represents an average of the two dCLNs from an individual mouse, n=4-9 mice/group per time point. Pooled data from three independent experiments. Scale: all images were taken at 10x. Images in (g) are zoomed insets of the 10x images. (i-k) Mice received TBI and then 2 hr after injury, fluorescently labeled anti-Lyve-1-488 antibodies (red) were injected i.c.m. into the CSF. Meningeal whole mounts were harvested 15 mins later and then stained for DAPI (blue) and Lyve-1-660 (gray). i) Representative images of meningeal whole mounts 2 hr after TBI stained for Lyve-1-488 (in vivo, red), Lyve-1-660 (ex vivo, gray) and DAPI. Solid box shows a zoomed inset of the hotspot along the transverse sinus on the right. Dashed box indicates the other hotspot not featured in the inset. Scale: left images were taken at 10x, the right images are zoomed insets of the 10x images. j) Percent area of Lyve-1-488 coverage at 2 hr post TBI. k) Distance traveled of Lyve-1-488 stain along transverse sinus 15 min after injection. Pooled data from two independent experiments (i-k). Each point represents an independent mouse and the error bars depict mean ± s.e.m. \**P* < 0.05, \*\**P* < 0.01, \*\*\**P* < 0.001, calculated by unpaired Students t-test (b,c,j,k), repeated-measures two-way ANOVA with Bonferroni’s post hoc test (d), and one-way ANOVA with Bonferroni’s post hoc test (h).

In order to assess whether meningeal lymphatic drainage is altered in this model of TBI, fluorescent beads were injected intra-cisterna magna (i.c.m.) at various timepoints after injury and the deep cervical lymph nodes (dCLN), meninges, and brain were harvested 2 hrs after injection to assess for the presence of beads (Figure 1e). Interestingly, when we examined cleared dCLN using confocal microscopy, we observed a substantial decrease in bead drainage to the dCLN in TBI mice (Figure 1f-h). As early as 2 hrs after injury, meningeal lymphatic function was severely impaired, as seen by decreased bead drainage into the dCLN (Figure 1f-h). Meningeal lymphatic drainage function remained significantly impaired out to at least two weeks post-injury (Figure 1f-h). Analysis of the brain and meningeal whole mounts revealed that the beads were taken up along the transverse sinuses as previously described [20], and were also detected around the fourth ventricle and in the cerebellum, both of which are areas close to the i.c.m. injection site (Supplementary Figure 2a,b). However, we were unable to detect any appreciable amount of beads in the systemic circulation even after TBI (Supplementary Figure 2c).

To determine whether the uptake of CSF into the meningeal lymphatic vasculature was altered in TBI, we examined the hotspots that exist along the transverse sinuses. These hotspots have been recently reported to be major areas of CSF uptake from the sub-arachnoid space [18, 23]. Fluorescently labeled Lyve-1 antibody was injected into the CSF i.c.m. at either 2 or 24 hrs after TBI, and then the meninges were harvested to examine uptake of CSF contents in the hotspots 15 mins after injection (Figure 1i-k and Supplementary Figure 3a-c). Analysis of the meningeal whole mounts revealed that there was significantly decreased uptake of fluorescently labeled Lyve-1 antibodies at the hotspots in mice that had received TBI 2 hrs prior when compared to mice that underwent a sham procedure (Figure 1i-k). Moreover, Lyve-1 antibody did not travel as far along the lymphatics lining the transverse sinus in the TBI mice at both 2 and 24 hrs post brain injury (Figure 1k and Supplementary Figure 3c). Taken together, these findings indicate that even mild forms of TBI can result in meningeal lymphatic dysfunction and that these deficits can persist out to at least 2 wks post-injury. Moreover, we find that disruption in CNS drainage function post-TBI is associated with impaired uptake of CSF at meningeal lymphatic hotspots.

### Increased intracranial pressure (ICP) results in meningeal lymphatic dysfunction

Elevated ICP is known to be a major driver of mortality after TBI and is associated with negative clinical outcomes [33, 34]. Because the lymphatic drainage deficit was substantial even 2 hrs after injury, we next examined whether there were associated changes in ICP after TBI. Two hrs after injury, TBI mice exhibited markedly increased ICP as compared to sham mice (Figure 2a,b). At later time points post-TBI, ICP levels stabilized slightly higher (∼5-7.5 mmHg) than baseline; however, the differences in ICP at these later timepoints are not found to be statistically significant (Figure 2a,b).

**Figure 2.**
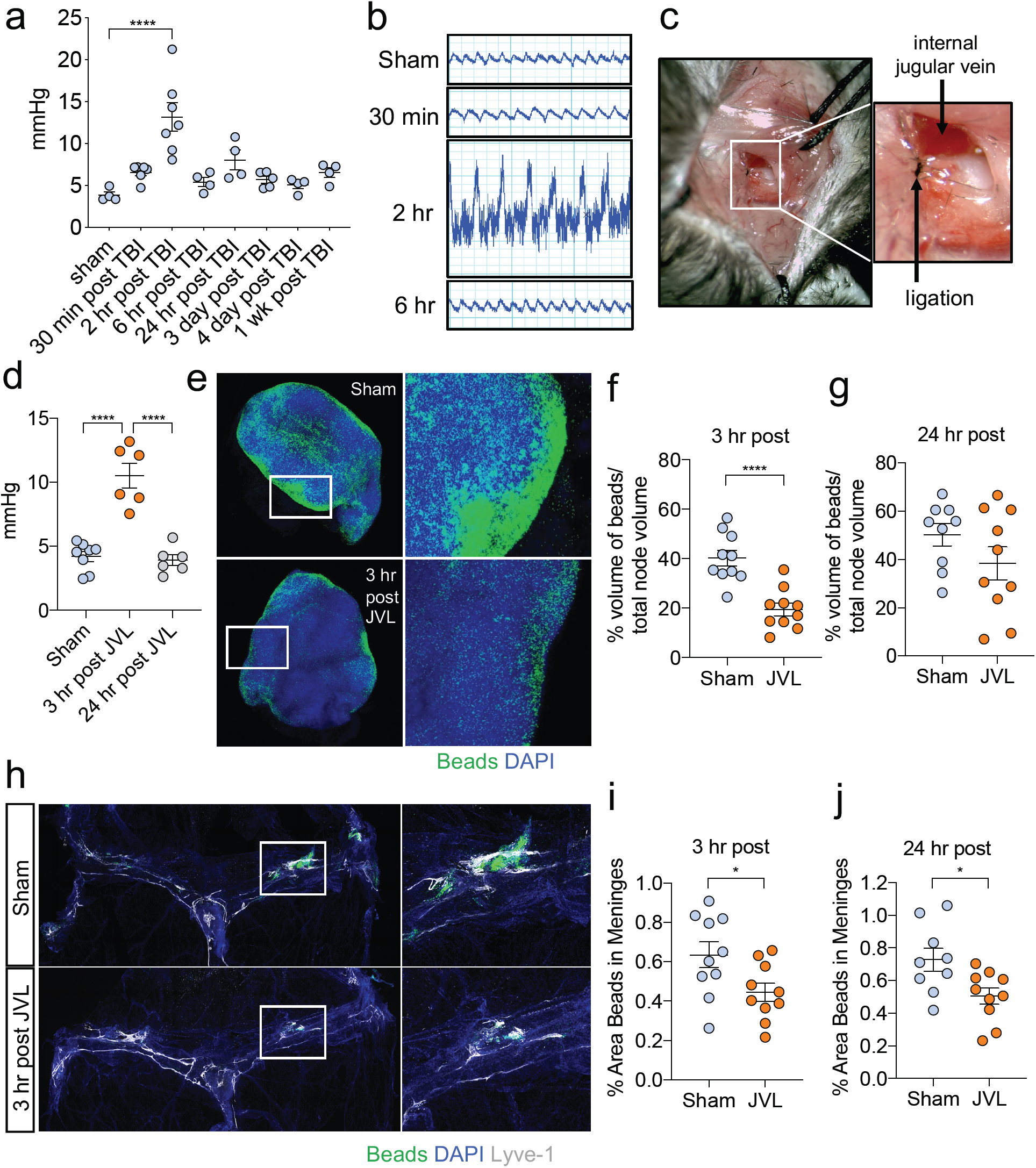
Increases in intracranial pressure disrupt CNS lymphatic drainage. a) Measurements of intracranial pressure (ICP) (b) and representative pressure readings were collected over an average of 6 mins at various timepoints after TBI. Pooled data from three independent experiments. c) Representative images of internal jugular vein ligation (JVL). d) ICP readings of mice that underwent bilateral jugular venous ligation or a sham procedure 3 or 24 hrs prior. Readings were collected over a span of 6 mins. Pooled data from three independent experiments. (e-j) The internal jugular vein was ligated bilaterally and then fluorescent beads were injected i.c.m. 3 hrs later. dCLN and meninges were then harvested from mice 2 hrs after bead injection. e) Representative images of dCLN and graph showing drainage of beads (f) 3 hrs and (g) 24 hr after jugular venous ligation. Each data point represents an average of the two dCLNs from an individual mouse. Pooled data from two independent experiments. Scale: all images were taken at 10x; the right images are zoomed insets of the 10x images. h) Representative images of meningeal whole-mounts stained with DAPI (blue) and Lyve-1-660 (grey) and graph depicting percent area of bead coverage (i) 3 hrs and (j) 24 hrs post-jugular vein ligation. Solid box shows a zoomed inset of the hotspot along the transverse sinus on the right. Scale: images were taken at 10x, the right images are zoomed insets of the 10x images. Pooled data from two independent experiments. Each point represents an independent mouse and the error bars depict mean ± s.e.m. \**P* < 0.05, \*\*\*\**P* < 0.0001, calculated by one-way ANOVA with Bonferroni’s post hoc test (a,d) and unpaired Students t-test (f,g,i,j)

Because the meningeal lymphatic vasculature is not associated with smooth muscle, it is especially vulnerable to changes in pressure and brain swelling inside the fixed skull [19, 22]. Therefore, we speculated that an acute rise in ICP might lead to disruptions in meningeal lymphatic drainage. To specifically test this, we subjected mice to bilateral internal jugular vein ligation (JVL, Figure 2c), which is known to increase ICP in both humans and mice [20, 35, 36]. Consistent with previous findings, we observed that jugular vein ligation substantially increased ICP to an average of 10 mmHg 3 hrs after surgical ligation, and that the ICP normalized by 24 hrs post ligation (Figure 2d) [20, 35, 36]. This reflected a similar acute spike in ICP that is seen after TBI (Figure 2a). To investigate what effect this rise in ICP has on meningeal lymphatic drainage function, we injected beads and assessed drainage to the dCLN at 3 and 24 hrs after jugular vein ligation. We found that there was significantly less drainage to the dCLN 3 hrs after bilateral internal jugular vein ligation (Figure 2e,f) and that there was reduced bead accumulation in the meninges of ligated mice (Figure 2h,i), indicating that there are deficits in the uptake of CSF contents in the CNS lymphatic vasculature. Moreover, we found that even after pressure normalized at 24 hrs (Figure 2d), there was still a prolonged period of decreased lymphatic drainage as seen by the diminished uptake of beads into the meningeal lymphatics (Figure 2j) and a trend towards decreased beads in the dCLN at 24 hrs post injury (Figure 2g). Collectively, these findings suggest that increased ICP is capable of provoking meningeal lymphatic dysfunction.

### Meningeal lymphatic dysfunction predisposes the brain to exacerbated neuroinflammation following TBI

TBI is an especially serious condition in the elderly and in individuals sustaining repetitive brain injuries [37-45]. For instance, similar injuries result in more severe pathology and neurological impairment in the elderly than in other age groups [37, 46]. Moreover, increasing evidence suggests that repetitive TBI can have devastating consequences that include CTE and mental disorders [39, 40, 47]. However, why TBI leads to worsened neurological disease in the elderly and following repetitive brain trauma remains poorly understood. Interestingly, it has recently been shown that CNS lymphatic drainage function significantly declines during aging [21, 48]. Moreover, as we demonstrated in Figure 1, a single head injury can provoke pronounced disruptions in meningeal lymphatic function. This led us to question whether pre-existing meningeal lymphatic dysfunction contributes to the exacerbated clinical disease following TBI, and if this might help to explain the increased severity of TBI disease seen in repetitive TBI and the elderly.

Therefore, to formally investigate how antecedent meningeal lymphatic deficits affect outcomes after TBI, we utilized a pharmacological approach to selectively ablate the meningeal lymphatic vessels before head injury. Visudyne, a photoconvertible drug that has been shown to effectively ablate lymphatic vasculature [20, 21, 28], was injected i.c.m. into the CSF and allowed to travel into the CNS lymphatics. A nonthermal 689-nm laser was then aimed through the skull to selectively photoablate the meningeal lymphatics. Mice were then rested for one week before receiving a brain injury (Figure 3a). Photoablation after Visudyne injection (Visudyne+laser) resulted in a significant decrease in area of the meninges covered by Lyve1-expressing lymphatic vessels in comparison to mice that received vehicle and laser treatment (Vehicle+laser) or Visudyne without laser treatment (Visudyne) (Figure 3b,c). Consistent with previous reports [20, 21, 28], the area of CD31^+^ blood vasculature was unchanged between all experimental groups (Figure 3b,d), indicating that the meningeal lymphatic vasculature was selectively ablated using this approach.

**Figure 3.**
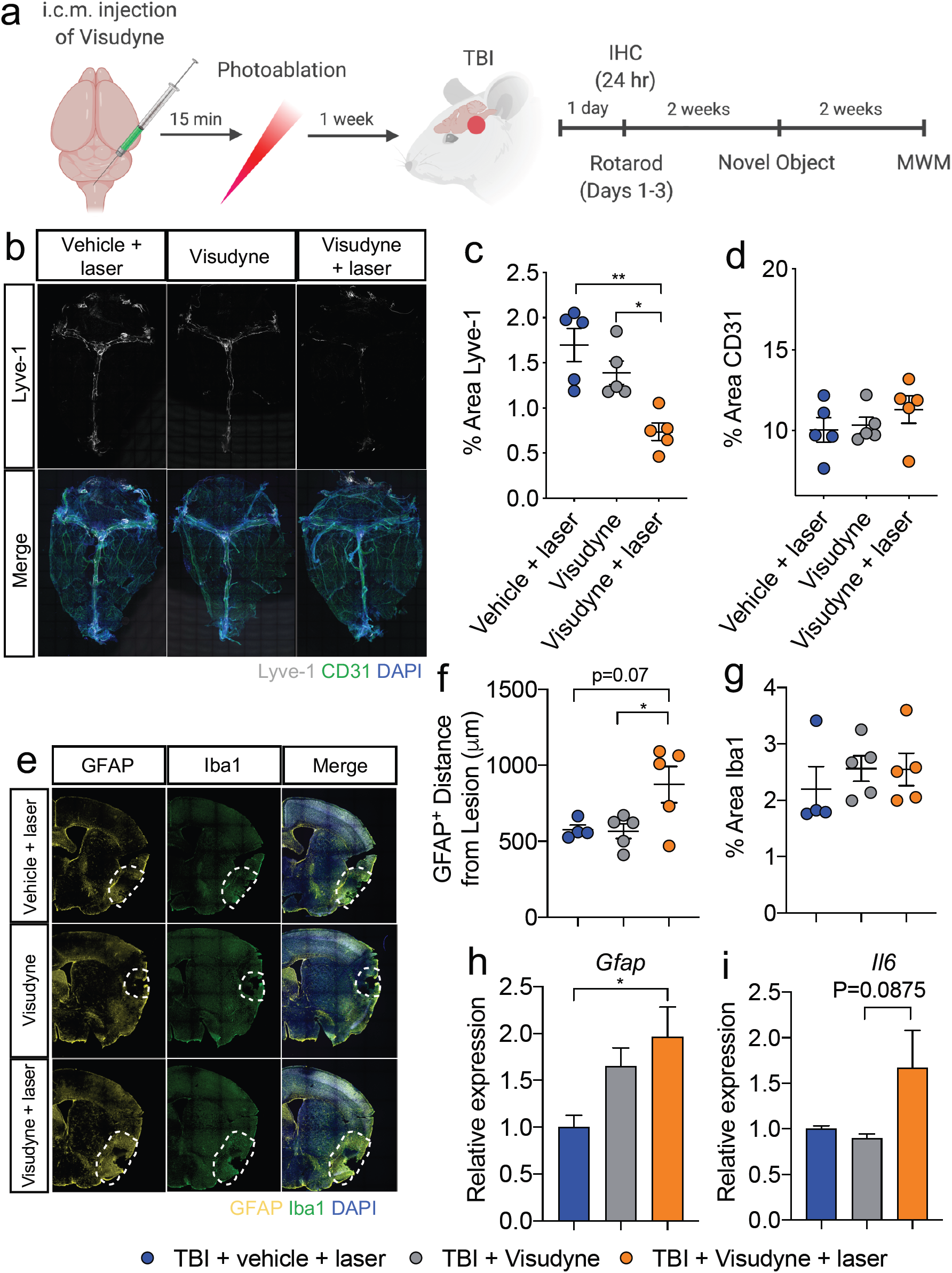
Pre-existing meningeal lymphatic dysfunction exacerbates neuroinflammation after TBI. a) Schematic of experimental layout. (b-i) Mice were subjected to an injection of Visudyne or vehicle i.c.m., and 15 mins later a red laser was directed at 5 spots along the sinuses through the skull. (e-i) After a week of recovery, mice received TBI or a sham procedure. b) Meningeal whole mounts were harvested 1 wk post-photoablation and then were stained with Lyve-1-660 (grey), CD31 (green), and DAPI (blue) to assess lymphatic and blood vasculature. Scale: all images were taken at 10x. Percent area coverage of (c) Lyve-1-660 or (d) CD31 in meningeal whole mounts. Representative data from 3 independent experiments. e-g) Brains from adult mice were harvested 24 hrs after TBI and representative images stained for Iba1 (green), GFAP (yellow) and DAPI (blue). Dashed lines indicate area of gliosis around lesion site. Scale: all images were taken at 10x. f) Distance of GFAP^+^ staining from the TBI lesion site. The average of 5 measurements around the lesion were calculated and averaged between two brain slices from each mouse. g) Percent area covered by Iba1 staining. h,i) RNA was extracted from the brains of mice 24 hrs post-TBI and expression of (h) *Gfap and* (i) *Il6* was evaluated by qPCR, n=6 mice/group. Each point represents an independent mouse and the error bars depict mean ± s.e.m. \**P* < 0.05, \*\**P* < 0.01, calculated by one-way ANOVA with Bonferroni’s post hoc test (c,d,f-i).

To explore how pre-existing meningeal lymphatic dysfunction influences neuroinflammation in TBI we first investigated changes in GFAP and Iba1 staining, as aggravated gliosis often correlates with worsened clinical outcomes in TBI [49-53]. We found that possessing defects in the meningeal lymphatic system before TBI (TBI+Visudyne+laser) results in greater distance covered by GFAP-expressing astrocytes surrounding the lesion site at 24 hrs post-injury (Figure 3e,f). We did not, however, observe any major differences in percent area covered by Iba1 staining between the groups 24 hrs after brain trauma (Figure 3e,g). Additionally, we observed increased expression of both *Gfap* and *Il6* expression in the TBI+Visudyne+laser group (Figure 3h,i). These results suggest that more severe neuroinflammation can unfold if meningeal lymphatic dysfunction already exists before TBI.

### Pre-existing meningeal lymphatic dysfunction before TBI results in more severe cognitive deficits

We were next interested in elucidating how pre-existing meningeal lymphatic dysfunction affects behavior and cognitive function following TBI. To explore this, we selectively ablated the meningeal lymphatics with Visudyne and photoconversion as described above, and then mice were subjected to either sham treatment or TBI one week later. We also included a cohort of mice that received Visudyne by i.c.m. injection but did not undergo photoablation (non-ablated). These non-ablated mice then received sham treatment or a TBI one week later. Taken together, this experimental setup provides controls for the surgical procedure, the laser photoablation, and the brain injury. All four groups of mice were then evaluated in behavioral tests to assess motor coordination and learning, cognitive function, and anxiety-related behaviors. Interestingly, while all four experimental groups exhibited similar performance on the accelerating rotarod at 24 hrs post-TBI, the mice possessing deficits in meningeal lymphatic function before TBI (ablated + TBI) showed impaired motor learning over days 2 and 3 of the accelerating rotarod test and consistently had a shorter latency to fall than the other control groups (Figure 4a). Indeed, the percent performance increase over three days in the ablated + TBI group was lower than any of the other control groups indicating that undergoing meningeal lymphatic photoablation before TBI results in impaired motor learning (Figure 4b). Moreover, mice that possessed meningeal lymphatic deficits before brain trauma exhibited impaired learning memory in the Morris water maze (MWM) test out to at least one month post-TBI. More specifically, the ablated + TBI group exhibited a decreased ability to learn over the training days as compared to mice that had received TBI alone (Figure 4c), and ablated + TBI mice also showed no improvement over the two days spent on the reversal task, whereas all other groups substantially improved in the reversal task (Figure 4d). The percent performance increase over the two reversal days showed that while all control groups improved, the group that had its meningeal lymphatic vessels ablated before TBI did not improve in its ability to find the platform over the two days (Figure 4e). Likewise, mice that underwent meningeal lymphatic photoablation before TBI also performed worse in the novel object recognition test (NORT) at two weeks post-brain injury (Supplementary Figure 4a,b). In comparison to the differences seen in cognitive function and learning, we observed similar performance between all experimental groups in behavioral tests to measure anxiety-related phenotypes (i.e. the open-field test and elevated plus maze) (Supplementary Figure 5a,b). Taken together, these results indicate that more severe cognitive deficits occur if CNS lymphatic dysfunction exists before brain injury. Moreover, by demonstrating that unresolved meningeal lymphatic dysfunction renders the brain more susceptible to worsened clinical outcomes after brain trauma, we provide insights into why TBI is especially devastating when the brain has not had sufficient time to recover between injuries.

**Figure 4.**
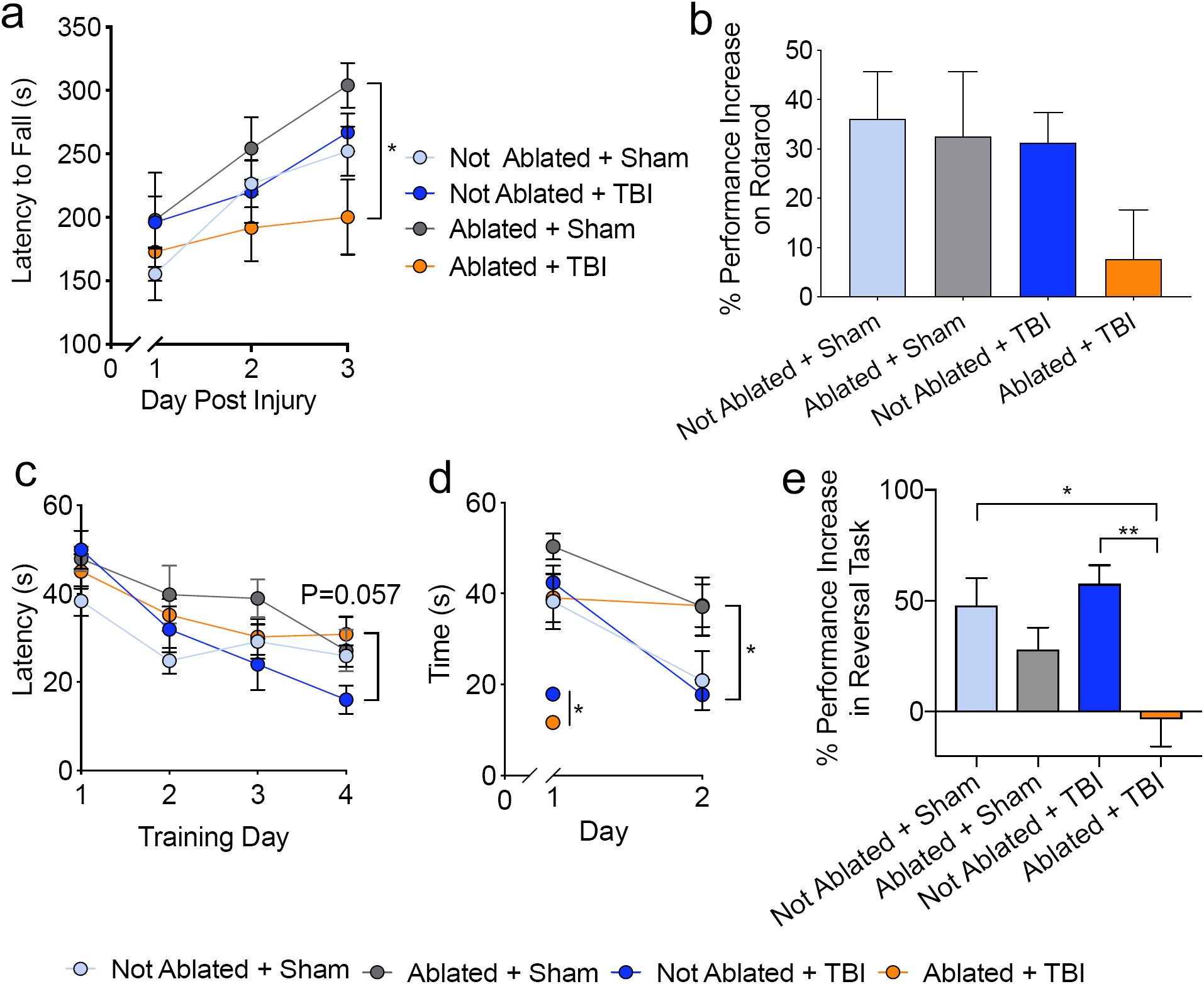
Pre-existing CNS lymphatic dysfunction before TBI results in impaired motor learning and memory. Mice were subjected to photoablation after Visudyne injection or control procedures and then to TBI or a sham procedure 1 week later. a,b) The accelerating rotarod test to assess motor function and motor learning was performed the first 3 days after TBI or sham procedure. a) Latency to fall over three days on the accelerating rotarod. Representative data from two independent experiments, n=6-7 mice/group. b) Percent performance increase on the rotarod. Pooled data from 2 independent experiments, n=6-16 mice/group. c-e) The Morris Water Maze (MWM) was performed 1 mo after injury to assess learning memory and spatial recognition. c) Latency to platform in the training and d) reversal days in the MWM was recorded, n=7 mice/group. e) Percent performance increase over the two reversal days in the MWM, n=7 mice/group. Each point represents an independent mouse and the error bars depict mean ± s.e.m. \**P* < 0.05, \*\**P* < 0.01, calculated by repeated-measures two-way ANOVA with Bonferroni’s post hoc test (a,e,f), and one-way ANOVA with Bonferroni’s post hoc test (b,g).

## DISCUSSION

Here, we report that meningeal lymphatic function is impaired after TBI and that this disruption begins almost immediately and can persist for weeks. We further show that ICP was significantly elevated at two hours post-brain injury, and that this increase in ICP can promote hotspot dysfunction in the uptake of CSF from the subarachnoid space. Moreover, we show that increased ICP is sufficient to cause meningeal lymphatic dysfunction. Our data also provide evidence that pre-existing lymphatic dysfunction, as may occur with repetitive TBI and aging, results in increased neuroinflammation and more severe cognitive deficits.

Several recent studies have shown that the meningeal lymphatic system is critical for modulating immune responses and inflammation in the CNS [20, 21]. Whether CNS lymphatic drainage is involved in promoting or resolving inflammation is likely specific to individual disease settings. For instance, mounting evidence indicates that lymphatic drainage plays a role in promoting autoimmunity by facilitating drainage of brain antigens to the peripheral dCLN. In the context of experimental autoimmune encephalomyelitis (EAE), ablation of the meningeal lymphatic vasculature was found to decrease disease severity through decreased CD4^+^ T cell infiltration to the spinal cord [20]. Consistent with a disease-promoting role for CNS lymphatic drainage in EAE, other studies have shown that lymphangiogenesis near the cribriform plate is a hallmark of disease progression and suggest that this may augment the peripheral immune response to myelin peptides [54]. In other instances, drainage of macromolecules and protein aggregates from the brain through the meningeal lymphatics is essential to maintain CNS health, as has recently been shown to be the case in mouse models of Alzheimer’s disease [21]. In this study, it was shown that blocking lymphatic drainage with Visudyne photoablation results in the accumulation of amyloid beta aggregates in the meninges and hippocampus.

Neuroinflammation and gliosis can often persist for months or even years post-brain trauma [8, 10, 53, 55]. The physiological processes involved in prolonging the inflammatory state of the TBI brain remain poorly defined. Based on our findings presented here, it is possible that impaired drainage of DAMPs such as amyloid beta, necrotic cells, and cellular debris from the brain could incite prolonged activation of the immune responses in the injured brain. Therefore, therapeutic interventions that promote functional recovery of the meningeal lymphatic system may offer novel strategies to help curtail the sustained neuroinflammation often seen following TBI.

Aggressive management of elevated ICP after a TBI may be one therapy that can limit lymphatic dysfunction if addressed immediately after injury. An acute rise in ICP after injury is a poor prognostic indicator in TBI patients, and is estimated to account for nearly half of all TBI mortalities [33, 34]. We found that the rise in ICP seen after TBI or jugular vein ligation is associated with decreased meningeal lymphatic drainage, indicating that changes in the CNS environment can rapidly impact lymphatic function. While acute rises in ICP after injury, caused by brain edema and swelling, may result in decreased drainage [33], how more prolonged rises in pressure affect the meningeal lymphatic vasculature and neuroinflammatory responses remain to be seen. Preventing a rapid and substantial rise in ICP after brain injury may allow for a more succinct immune response due to more efficient drainage of DAMPs from the CNS.

Because clearance of interstitial fluid (ISF) containing solutes and macromolecules from the brain parenchyma relies on perivascular routes, termed the glymphatic system [56-59], we anticipate that changes in ICP would also affect glymphatic function. Indeed, our findings are consistent with the data that show the glymphatic system to be impaired after TBI [57, 60]. The glymphatic system and lymphatic system are inherently linked [56, 61]. The glymphatic system is responsible for transport of ISF to the CSF surrounding the brain, and the lymphatic system takes up CSF/ISF for transport into the periphery [57, 59, 61, 62]. While decreased ISF transport from the parenchyma to the sub-arachnoid space may lead to decreased lymphatic drainage [57, 58], it has also been shown that impaired lymphatic drainage decreases recirculation of macromolecules through the glymphatic route [21]. Our findings indicate that, in addition to the previously described glymphatic dysfunction, there is also lymphatic dysfunction after TBI. This was shown by directly targeting the meningeal lymphatic vasculature through pharmacologic photoablation. In these studies, we found that lymphatic dysfunction results in worsened behavioral outcomes and increased neuroinflammation. Additionally, injections into the cisterna magna, as was done in our studies to assess lymphatic drainage efficiency, largely bypass the need for glymphatic clearance, as the meningeal lymphatic network takes up CSF directly from this compartment. Therefore, the combination of impairment in these two systems likely coalesce to decrease clearance of CNS-derived toxins, protein aggregates, and macromolecules generated following brain trauma. Preventing the rapid rise in ICP seen after TBI may provide a route in which to address both the glymphatic and lymphatic dysfunction that persists after injury.

Interestingly, TBI has been strongly linked to increased risks of developing numerous other neurological disorders later in life including CTE, Alzheimer’s disease, amyotrophic lateral sclerosis, and multiple mental disorders [2-7]. Our understanding of how brain trauma contributes to the development of these other neurological disorders at the mechanistic level is currently limited. Like TBI, the majority of these CNS disorders are also characterized by neuroinflammation and impaired clearance of DAMPs (e.g. protein aggregates and neurotoxic debris). One could envision that TBI-induced disruptions in meningeal lymphatic function and the buildup of DAMPs in the brain could set off a series of events that ultimately lead to other forms of neurological disease down the road, although future studies are needed to formally test this hypothesis. Investigation of whether rejuvenation of meningeal lymphatic function after TBI is capable of limiting the risk of disease sequelae later in life should help to address this in future work.

For reasons that remain poorly understood, sustaining a second head injury before the brain has recuperated from prior head trauma can have devastating consequences. Furthermore, it has been shown that repetitive TBIs result in more serious long-term outcomes when compared to a single TBI [44, 45, 47, 63, 64]. Indeed, recent reports in the scientific literature and media have highlighted several high-profile cases of repetitive TBI and its devastating consequences that include CTE and suicide [39, 40, 47, 65]. Improved understanding of what makes the injured brain more vulnerable to more severe pathology and neurological demise following secondary head trauma will lead to improved treatment practices. Our findings presented in this paper suggest that the disruptions in meningeal lymphatic function that occur following TBI, may contribute at some level to the more severe neuroinflammation and cognitive deficits commonly observed in repetitive TBI. Indeed, we show that a single mild brain injury does not result in gross behavioral disturbances alone, but the second ‘hit’ to an already compromised lymphatic system is enough to result in pronounced behavioral deficits. The lack of defined guidelines for when individuals can safely return to high-risk activities following TBI is a significant problem for caregivers. Our work suggests that evaluating meningeal lymphatic drainage recovery post-injury might provide clinicians with a much-needed empirical test to inform athletes and military personnel when it is safe to return to action. However, improved diagnostics must first be developed to accurately measure meningeal lymphatic function in humans for this to come into practice in the future.

TBI is an especially serious threat to health in the elderly, where it is a leading cause of death and disability [41-43, 46, 66]. Even though the elderly only account for 10% of all TBI cases, over 50% of all TBI-related death occurs in individuals over the age of 65 [67]. It has been extensively shown that similar injuries result in more severe pathology and neurological impairment in the elderly than in other age groups [37, 46]; however, the cause of t.R.L. performed experiments. A.C.B. and J.R.his is currently not well understood. Interestingly, several recent studies have shown that the meningeal lymphatics are impaired with aging [21, 23, 48]. Moreover, it was shown that boosting lymphatic function by widening the vessel diameter can decrease the cognitive deficits seen in aged mice [20]. In our studies presented here, we show that photoablation of meningeal lymphatic vessels before head injury can predispose the brain to exacerbated clinical outcomes following TBI. Therefore, it is feasible that aging-associated deterioration of CNS lymphatic function may contribute at some level to the especially devastating consequences of TBI in the elderly. Although, future studies aimed at uncovering how the meningeal lymphatic system influences TBI disease progression in the elderly are needed to more fully address this.

Overall, the work described here provides new insights into how the meningeal lymphatic system is impacted by TBI and also how pre-existing defects in this drainage system can predispose the brain to exacerbated neuroinflammation and cognitive deficits following brain injury. We show that even mild forms of TBI can result in pronounced defects in meningeal lymphatic function that can last out to at least two weeks post-injury. Mechanistically, we show that closed-skull TBI is associated with elevated ICP and that this can directly contribute to disruptions in meningeal lymphatic drainage function. Continued efforts to better characterize the role of the meningeal lymphatic system in brain injury may help to identify improved strategies to treat TBI and/or limit trauma-associated disease sequelae.

## Abbreviations

CNS: central nervous system
CSF: cerebrospinal fluid
CTE: chronic traumatic encephalopathy
DAMPs: danger/damage-associated molecular patterns
dCLN: deep cervical lymph nodes
hpi: hours post-injury
ICP: intracranial pressure
i.c.m.: intra-cisterna magna
IHC: immunohistochemistry
ISF: interstitial fluid
JVL: jugular vein ligation
NORT: novel object recognition test
MWM: Morris water maze
TBI: traumatic brain injury
wpi: weeks post-injury

## ACKNOWLEDGEMENTS

We thank members of the Lukens lab and the Center for Brain Immunology and Glia (BIG) for valuable discussions. This work was supported by The National Institutes of Health/National Institute of Neurological Disorders and Stroke (R01NS106383; awarded to J.R.L.), The Alzheimer’s Association (AARG-18-566113; awarded to J.R.L.), The Owens Family Foundation (Awarded to J.R.L.), and The University of Virginia Research and Development Award (Awarded to J.R.L.). A.C.B. and M.A.K. were supported by a Medical Scientist Training Program Grant (5T32GM007267-38) and an Immunology Training Grant (5T32AI007496-25). H.E.E was supported by a Cell and Molecular Biology Training Grant (T32GM008136). C.R.L was supported by a NIH National Institute of General Medical Sciences predoctoral training grant (3T32GM008328) and a Wagner Fellowship. E.L.F. was supported by a National Multiple Sclerosis Foundation Postdoctoral Fellowship (FG-1707-28590).

## Contributions

A.C.B, J.K. and J.R.L. designed the study; A.C.B., M.E.H., I.S., E.L.F., C.R.L., C.A.M., N.N., M.A.K. and J.R.L. performed experiments. A.C.B. and J.R.L. analyzed data and wrote the manuscript; J.R.L. oversaw the project.

