## Supplementary figures and legends for "Meningeal lymphatic dysfunction exacerbates traumatic brain injury pathogenesis"

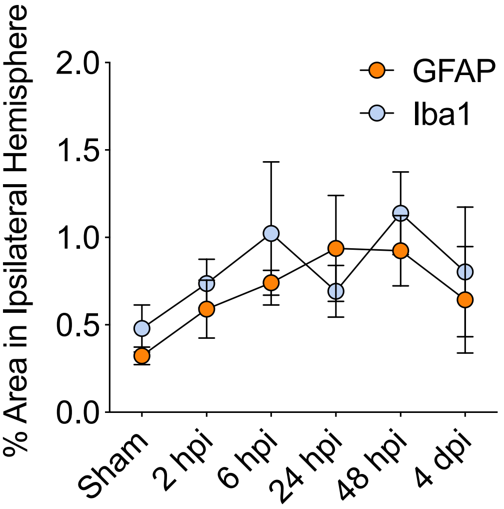


Supplementary Figure 1

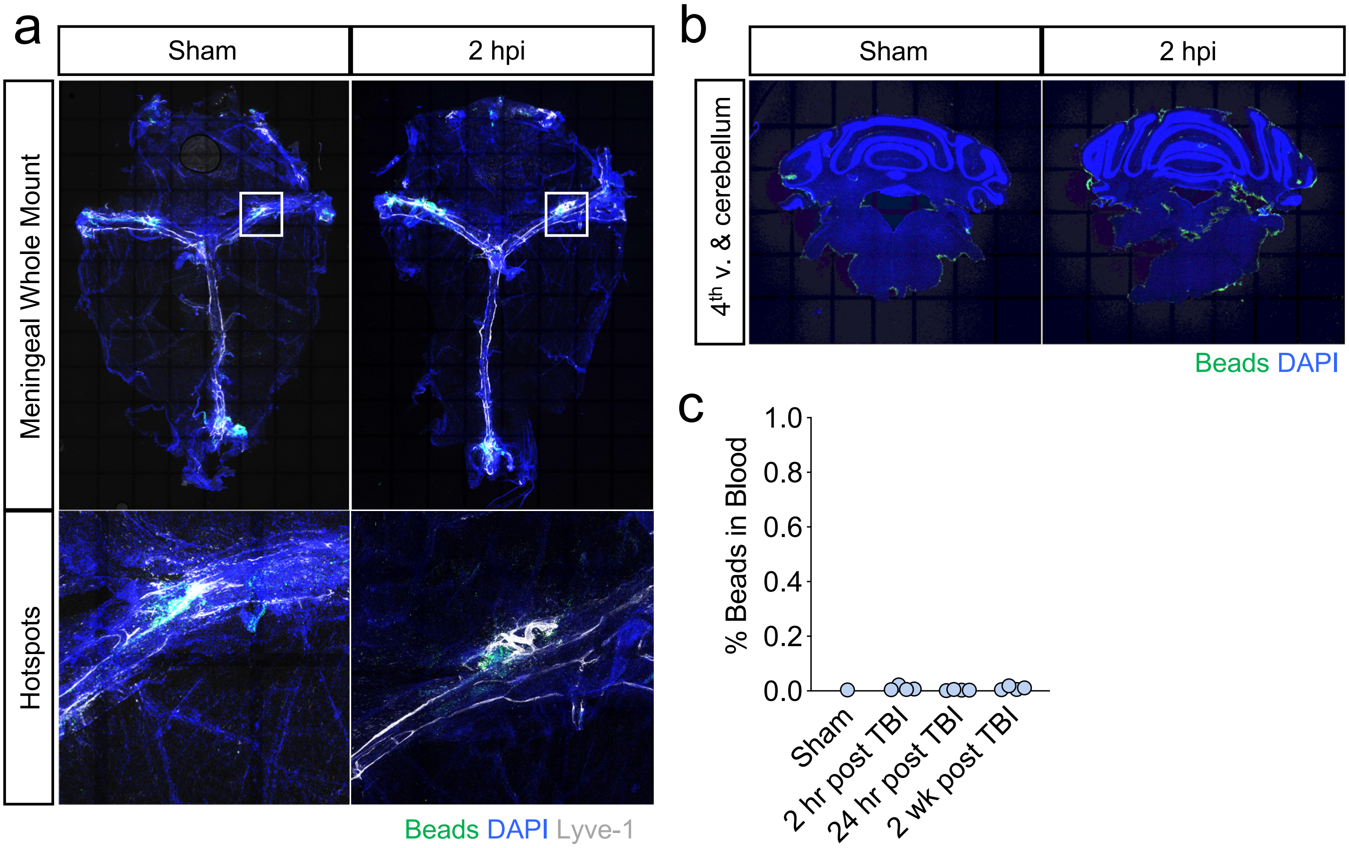


Supplementary Figure 2

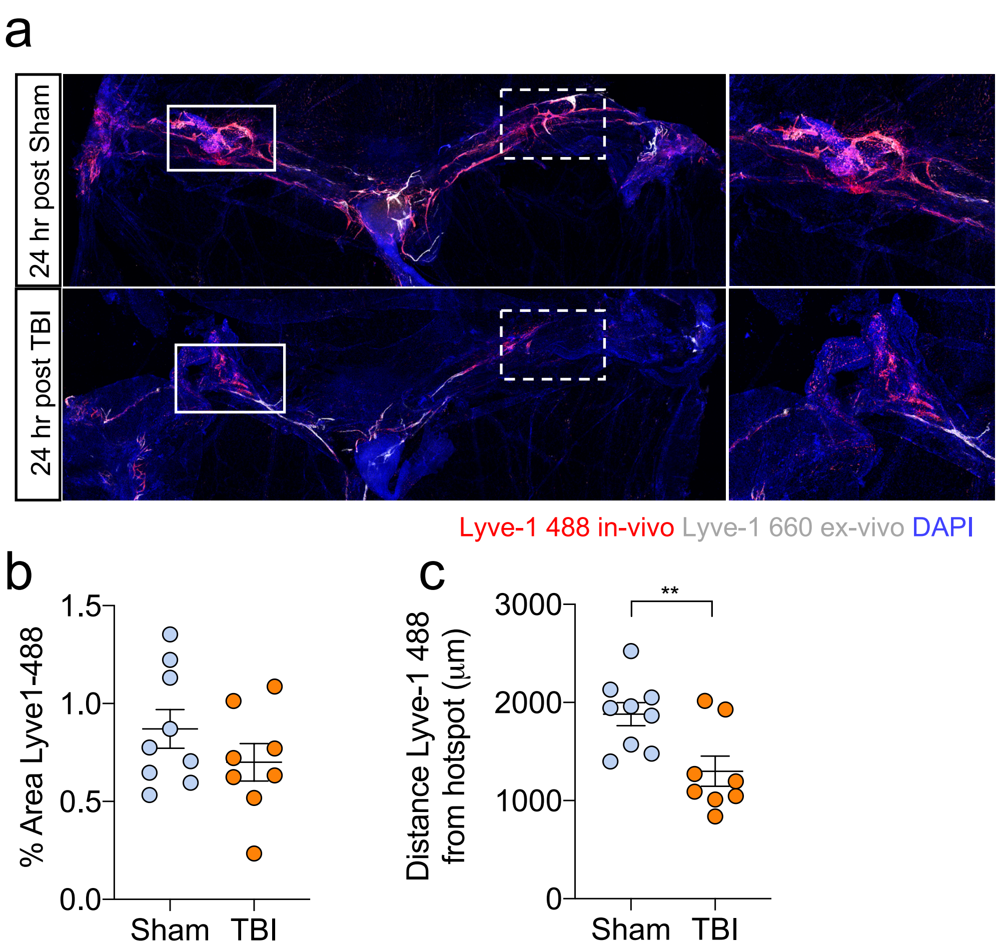


Supplementary Figure 3

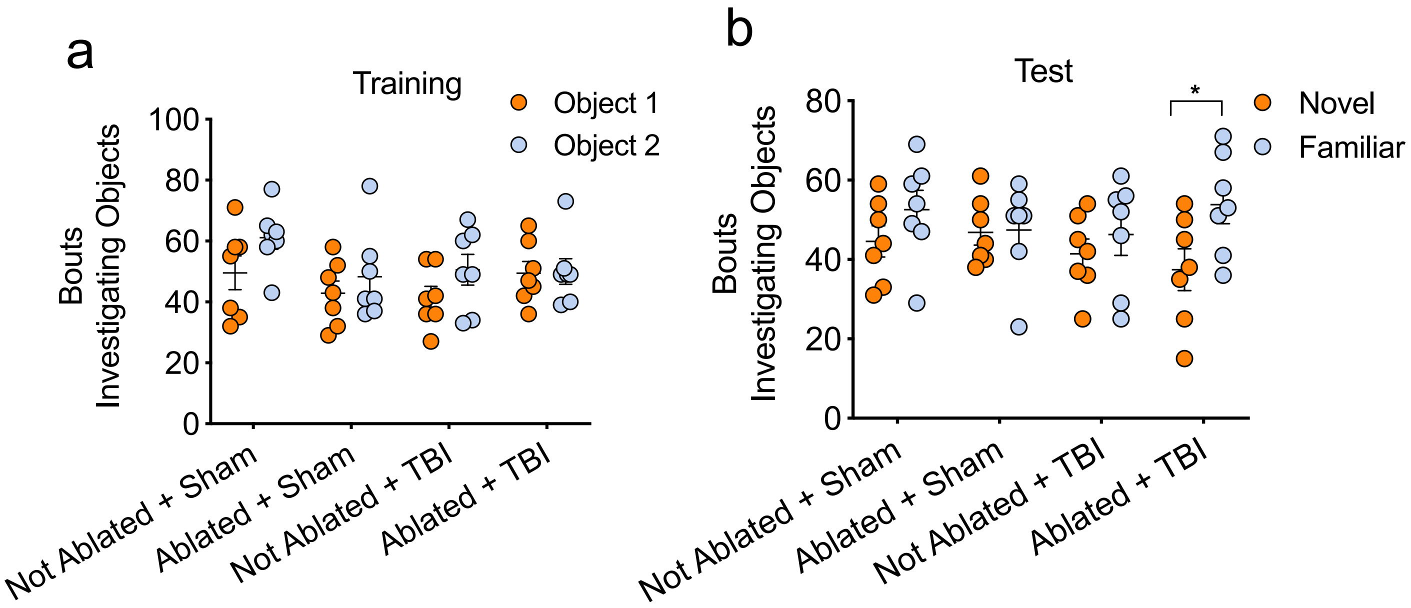


Supplementary Figure 4

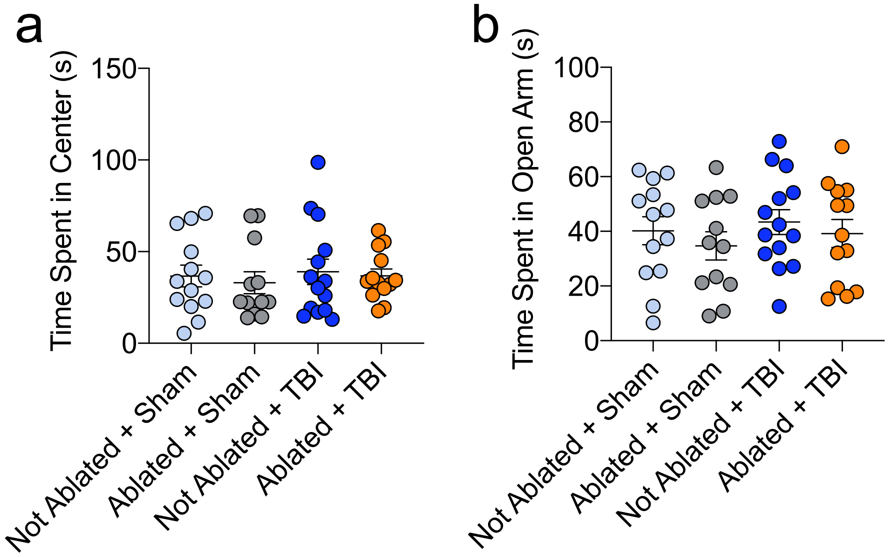


Supplementary Figure 5

### SUPPLEMENTARY FIGURE LEGENDS

**Supplementary Figure 1. Mild closed-skull TBI results in modest increases in gliosis**. Brains were harvested at various timepoints after TBI and stained for GFAP and Iba1. Percent area coverage of Iba1 and GFAP in the brain hemisphere containing the TBI lesion site, n=3-7 mice/group per timepoint. Pooled data from 2 independent experiments. All data are presented as mean ± s.e.m.

**Supplementary Figure 2. Beads are found in the meninges, fourth ventricle, and cerebellum but not the blood after i.c.m. injection**. Mice received TBI or sham treatment and then received 0.5 μm fluorescent beads (green) by i.c.m. injection 2 hrs later. Meningeal whole mounts and brains were harvested 2 hr after injection. a) Representative images of meningeal whole mounts stained with DAPI (blue) and Lyve-1-660 (gray) after TBI or a sham procedure and fluorescent bead injection. Solid box shows inset of bead uptake at the hotspot along the transverse sinus, Scale: all images were taken at 10x, the images below are zoomed insets of the 10x images. b) Representative images of fluorescent beads in the fourth ventricle and cerebellum. c) Frequency of beads in the blood 2 hrs, 24 hrs, or 2 wk post TBI. Each point represents an independent mouse and the error bars depict mean ± s.e.m.

**Supplementary Figure 3. Meningeal lymphatic uptake of CSF at hotspots is impaired after TBI**. a-c) Mice received TBI or sham treatment and then fluorescently labeled anti-Lyve-1-488 antibodies (red) were injected i.c.m. into the CSF 24 hrs later. Meningeal whole mounts were then harvested and ex vivo stained with anti-Lyve-1-660 (gray) 15 mins following injection with Lyve-1-488 antibodies. a) Representative images of meningeal whole mounts 24 hrs after TBI stained for Lyve-1-488 (in vivo, red), Lyve-1-660 (ex vivo, gray) and DAPI (blue). Solid boxes show zoomed insets of the hotspots along the transverse sinus on the right. Dashed boxes indicate the other hotspots not featured in the insets. Scale: left images were taken at 10x, the right images are zoomed insets of the 10x images. b) Percent area of Lyve-1-488 coverage at 24 hrs post-TBI. Pooled data from two independent experiments. c) Distance traveled of Lyve-1-488 stain along transverse sinus 15 min after injection. Pooled data from two independent experiments. Each point represents an independent mouse and the error bars depict mean ± s.e.m. $^{**}P < 0.01$, calculated by Student's t-test.

**Supplementary Figure 4. Pre-existing lymphatic dysfunction prior to TBI results in abnormalities in the novel object recognition test.** a,b) The Novel Objection Recognition Test (NORT) was performed 2 weeks after TBI to assess memory. a) Time spent investigating two identical objects was recorded for each mouse on the training day of the NORT. Object 1 and object 2 represent two identical objects. b) Time spent investigating the familiar and novel object was recorded on the test day of the NORT where the familiar object is the same object used in the training phase, and the novel object is a new object with a different texture, shape and color. Each point represents an independent mouse and the error bars depict mean ± s.e.m. **P* < 0.05, calculated by paired Students t-tests between the Object 1/Object 2 pairs or Novel/Familiar object pairs (a,b).

**Supplementary Figure 5. Deficits in meningeal lymphatic function prior to TBI do not significantly impact performance in the open-field test or the elevated plus maze**. Mice were subjected to photoablation after Visudyne injection or control procedures and then to TBI or a sham procedure one week later. a) The open-field test was performed 2 weeks after injury to measure anxiety-like behaviors. The time spent in the center was evaluated. Pooled data from two independent experiments. b) Time spent in the open arm of the elevated plus maze (EPM) was measured to further assess anxiety-like behaviors. Pooled data from two independent experiments. Each point represents an independent mouse and the error bars depict mean ± s.e.m. No statistical differences were observed.
